# Increasing NPYergic transmission in the hippocampus rescues aging-related deficits of long-term potentiation in the mouse dentate gyrus

**DOI:** 10.1101/2023.08.11.552947

**Authors:** Katharina Klinger, Miguel del Ángel, Gürsel Çalışkan, Oliver Stork

## Abstract

Loss of neuropeptide Y (NPY)-expressing interneurons in the hippocampus and decaying cholinergic neuromodulation are thought to contribute to impaired cognitive function during aging. However, the interaction of these two neuromodulatory systems in maintaining hippocampal synaptic plasticity during healthy aging has not been explored so far. Here we report profound sex differences in the Neuropeptide-Y (NPY) levels in the dorsal dentate gyrus (DG) with higher NPY concentrations in the male mice compared to their female counterparts and a reduction of NPY levels during aging specifically in males. This change in aged males is accompanied by a deficit in theta burst-induced long-term potentiation (LTP) in the medial perforant path-to-dorsal DG (MPP-DG) synapse, which can be rescued by enhancing cholinergic activation with the acetylcholine esterase blocker, physostigmine. Importantly, NPYergic transmission is required for this rescue of LTP. Moreover, exogenous NPY application alone is sufficient to recover LTP induction in aged male mice, even in the absence of the cholinergic stimulator. Together, our results suggest that in male mice NPYergic neurotransmission is a critical factor for maintaining dorsal DG LTP during aging.

## 1. INTRODUCTION

Neuronal hyperactivity and reduced capacity of synapses to express long-term plasticity are commonly observed in neurodegenerative diseases ^1,2^ and during aging ^3,4^. The hippocampus, one of the hub regions that play a central role in learning and memory ^5^, shows such hyperactivity and aberrant synaptic plasticity ^3,4^. Therefore, studies that aim at identifying treatment strategies for normalizing hippocampal function are of utmost importance to mitigate cognitive decline during aging.

Neuropeptide-Y (NPY) belongs to the pancreatic polypeptides class and is widely distributed in the nervous system ^6^. NPY is highly expressed in the hilus of the dentate gyrus (DG) ^7,8^, the gate of the hippocampal trisynaptic circuit that plays a pivotal role in encoding contextual and spatial information ^7,9^. The DG receives excitatory perforant path (PP) projections from the entorhinal cortex that are substantially affected in the aging process in rodents ^10^ and humans ^11^. We and others have previously shown that NPY plays a crucial role in controlling DG circuit function and PP-induced physiological responses in young adult mice ^7,12^. Importantly, a substantial decline in the levels of NPY and NPY receptor type 1 (Y1-R), which is highly expressed in DG excitatory granule cells (GC), has been shown during the aging process ^13,14^. However, to the best of our knowledge, no studies have yet systematically investigated the mechanistic involvement of NPYergic transmission in the PP-induced neurotransmission and plasticity in the DG of aged mice.

Disturbances of the hippocampal circuit function during aging are accompanied by aberrant functional or structural alterations in the long-range cholinergic projections (e.g., synaptic loss, cholinergic axonal degeneration) from the medial septum to the hippocampus ^15,16^ and are frequently used to model age–related changes in hippocampal function ^17^. Accordingly, the beneficial effects of acetylcholine esterase (AChE) inhibitors for the treatment of Alzheimer’s Disease and on long-term plasticity in aged rodents are well documented ^18,19^. It is important to note that cholinergic modulation of NPYergic neurotransmission is critically involved in the modulation of DG circuit function and DG-dependent behaviors ^7,20^. Specifically, enhanced cholinergic drive leads to Y1-R-dependent NPYergic signaling and altered DG output via activation of the hilar Somatostatin (SST) containing interneurons ^7^, most of which co-express NPY ^8,21^. Intriguingly, to date, the potential use of such synergistic interactions between the cholinergic and NPYergic systems in sustaining hippocampal DG neurotransmission and plasticity during aging has not been systematically explored.

Therefore, we addressed the potential involvement of NPY in mediating cholinergic modulation of plasticity in the aging hippocampus and explored the possibility that enhancement of NPYergic signaling may provide a resource for alleviating age-induced synaptic transmission and plasticity changes. It needs to be considered that both the cholinergic and the NPYergic system display remarkable sex differences. Particularly, the reduction in the density of cholinergic fibers targeting the DG is significantly larger in aged male rats than in aged females ^22^. Moreover, NPY levels differ between sexes in the hippocampus, with higher NPY levels in young males compared to female counterparts ^23^. Based on the decay of NPY expression that we observed specifically in the dorsal DG of aged (>20-month-old) male mice, we collected evidence that NPYergic signaling is required for the cholinergic (via physostigmine) alleviation of aging-related LTP deficits and that exogenous NPY application is sufficient to ameliorate the LTP deficit in these animals.

## 2. METHODS

### 2.1. Animals

Wild-type male and female C57BL/6 mice (C57BL/6BomTac; M&B Taconic, Germany) were housed and bred in the animal facility of the Department of Genetics and Molecular Neurobiology, Institute of Biology, Otto von Guericke University Magdeburg under standard laboratory conditions. Young-adult (2-4 months) and aged (>20 months) males and females were kept in groups of two to five individuals in an inverse 12 h light/ dark cycle (lights on from 7 PM – 7 AM with a 30 min dawn phase). They had access to food and water *ad libitum*. All experiment preparations were conducted during the animals’ dark (active) phase, between 8 AM and 3 PM.

### 2.2. Electrophysiology

#### 2.2.1. Slice Preparation

To investigate synaptic plasticity and transmission during aging and under different pharmacological conditions, young-adult and aged mice of both sexes were firstly deeply anesthetized with isoflurane and then decapitated. Brains were rapidly (within ∼60 s) removed and placed into cold (4–8 °C) carbonated (5% CO2 / 95% O2) artificial cerebrospinal fluid (aCSF) containing (in mM) 129 NaCl, 21 NaHCO3, 3 KCl, 1.6 CaCl2, 1.8 MgSO4, 1.25 NaH2PO4, and 10 glucose. Parasagittal slices containing the dorsal hippocampus were obtained by cutting the brain at an angle of about 12° on an angled platform. The four most dorsal slices were transferred to an interface chamber perfused with aCSF at 32 ± 0.5 °C (flow rate: 1.8ml ± 0.2 ml per min, pH 7.4, osmolarity ∼ 300 mosmol kg−1). After cutting, the slices were left to rest for 1 hour before starting the recordings.

#### 2.2.2. Field potential recordings

For dorsal DG electrophysiology, one glass electrode, filled with aCSF (∼ 1 MΩ), was placed at 70–100 μm depth into the mid-molecular cell layer (ML). The stimulation of the medial PP (MPP) was performed with a bipolar tungsten wire electrode, with exposed tips of ∼ 20 μm and tip separations of ∼ 75 μm (electrode resistance in aCSF: ∼ 0.1 MΩ, world precision instruments (WPI); Friedberg, Germany). A second bipolar tungsten electrode (same parameters as in the MPP) was placed into the hilus of the dorsal DG to elicit an antidromic stimulation during LTP induction. An input-output (I-O) curve for each stimulation electrode was recorded after stabilization of the responses (0.033 Hz, pulse duration: 100 μs) over 20 min. The baseline excitability and maximal synaptic response were measured by obtaining the I-O curve using pulses with stimulation intensities ranging from 10 to 75 μA. The stimulus intensity that resulted in ∼ 50% of the maximum field excitatory post-synaptic potentials (fEPSP) amplitude of the ML after orthodromic stimulation was subsequently used for baseline recordings with orthodromic stimulation. While the stimulus intensity that resulted in ∼ 70% of the maximum fEPSP amplitude of the ML after antidromic stimulation was used for the antidromic stimulation during LTP induction. The appropriate placement of the orthodromic stimulation was verified through paired-pulse stimulation with different inter-pulse intervals right after obtaining the Input-output (I-O) curve before starting the baseline recording. The characteristic feature of the MPP-DG synapse is the consistent paired-pulse depression at 50 ms inter-pulse interval ^4,24^. In all protocols, baseline and post-theta-burst stimulation (TBS) recordings were done with orthodromic stimulation (in the MPP). Exclusively, the four repetitions repeated at 0.1 Hz of TBS (one burst consisting of 10 pulses at 100Hz-repeated 10 times at 5 Hz) ^25^ were done with orthodromic followed by antidromic stimulation (in the hilus of the DG). Signals were pre-amplified using a custom-made or EXT20-F amplifier (npi electronics, Tamm, Germany) and low-pass filtered at 3 kHz. Signals were sampled at a frequency of 10 kHz and stored on a computer hard disc for offline analysis. The fEPSP slopes (20-80%) were analyzed offline using self-written MATLAB-based analysis tools (MathWorks, Natick, MA, USA).

#### 2.2.3. Pharmacological interventions during electrophysiological recordings

The effect of selective Y1-R blockade and NPY on the fEPSP slope at the MPP-DG synapse during neurotransmission and LTP responses in young and aged males were evaluated in two different experiments. Either the selective Y1-R antagonist BIBP3226 (1 μM; Cat.-No. 2707; Tocris, Bristol, UK) or NPY (1 μM; Cat.-No. 90880-35-6; Cayman Chemicals, Ann Arbor, Michigan, USA) was added to the aCSF for 20 min or 40 min, respectively, (until TBS) after 20 min baseline recording. To evaluate the effect of constitutive cholinergic activation and selective Y1-R blockade under such cholinergic activation, the AChE inhibitor physostigmine hemisulfate (PHY; 2 μM; Cat.-No. sc-203661; Santa Cruz Biotechnology, Dallas, Texas, USA) or PHY combined with BIBP3226 (1 μM) was perfused in slices of young and aged male mice. PHY was applied for 60 min or 40 min, followed by 20 min BIBP3226 (1 μM) application after an initial 20 min baseline recording. In all experiments, the perfusion solution was changed back to aCSF at the time point of TBS.

### 2.3. Protein analysis

Mice were deeply anesthetized with isoflurane and killed by cervical dislocation. After cervical dislocation, the mice were decapitated, and the brains were carefully removed from the skull and transferred to ice-cold phosphate buffer saline. For protein analysis, the dorsal DG was dissected manually ^26^. Samples were snap-frozen with liquid nitrogen and stored at -80°C.

#### 2.3.1. Western Blot (WB)

Hippocampal tissue (dorsal DG) of aged and young males and females were mechanically homogenized on ice in cold Laurylmaltoside-buffer containing 1% Laurylmaltoside, 1% NP-40, 1 mM Na_3_VO_4_, 2 mM EDTA, 50 mM Tris-HCl pH 8.0, 150 mM NaCl, 0.5% deoxycholate, 1 mM NaF 1mM AEBSF protease inhibitor (cat. No. 78431; Thermo Fisher Scientific, Massachusetts, USA), 1 µM Pepstatin A (cat. No. 78436; Thermo Fisher Scientific, Massachusetts, USA), and 1 Tablet of Pierce protease inhibitor (cat. No. A32963; Thermo Fisher Scientific, Massachusetts, USA). The protein concentration was quantified with the RC DC Protein assay Kit II (cat. No. 5000122; Bio-Rad Labortaories Inc, California, USA). After protein concentration assessment, the samples were prepared for immunoblot analysis by adding sample buffer (10% glycerol, 60 mM Tris HCL pH 6.8, 2% SDS, 0.01% Bromophenol blue, and 1,25% mercaptoethanol). For electrophoretic separation, an SDS Bis-Acrylamide gel was loaded with 20 µg of protein from each sample and a standard protein marker (PageRuler Plus, cat. No. 11852124; Thermo Fisher Scientific, USA). The proteins were transferred to FL-PVDF membranes (cat. No. IPFL00010; Merck Millipore; Massachusetts, USA). The membranes were incubated with ms anti-Y1-R (1:500; Santa Cruz Biotechnology, Texas, USA, cat. No. #sc-393192) primary antibody solution at 4°C overnight. A near-infrared labeled secondary antibody IRDye 800CW or IRDye680CW (1:10,000; LI-COR Biosciences, Nebraska, USA) was used to detect the primary antibody. Finally, the imaging of the antibodies was performed using the Odyssey Imaging system (LI-COR Biosciences, Nebraska, USA), and the protein signal was quantified in the Image Studio software (LI-COR Biosciences, Nebraska, USA).

#### 2.3.2. Enzyme-linked immunosorbent assay (ELISA)

To determine the NPY peptide levels, 5 µl of the protein extract from each sample was prepared in a final concentration of 1 µg/ µl and analyzed with the ELISA kit for NPY (Product No. CEA879Hu; UOM: 96T; with a minimum detectable dose less than 9.44 pg/mL) according to manufacturer’s instructions (cloude-clone corp., Texas, USA).

### 2.4. Statistics

Data are reported as mean ± standard error of the mean (SEM). Before the statistical comparison of electrophysiological data, outliers were identified after ROUT, and normality was evaluated with the D’Agostino-Pearson test for all statistical tests. Statistical analysis of I-O and FV curves was performed using two-way repeated measure analysis of variance (ANOVA) with Geisser-Greenhouse correction followed by post-hoc comparison using Fisher’s least significant difference (LSD) method. Baseline transmission data was normalized to aCSF condition (10 min before application of the drug of interest), whereas LTP data were normalized to the data obtained 10 min prior to TBS. An in-slice comparison using either a one-tailed paired t-test or Wilcoxon signed-rank test was performed to determine successful LTP induction. Furthermore, to evaluate statistical differences in the LTP levels or pharmacological treatment effects across different groups, either a two-tailed unpaired t-test, Mann-Whitney U test, or two-way ANOVA, where appropriate, was performed. ELISA and WB data were analyzed with a two-way ANOVA Geisser-Greenhouse correction and Fisher’s LSD post-hoc test. Graphs and statistical tests were conducted with GraphPad Prism (version 9.4.1(681); Dotmatics, Boston, Massachusetts, USA). Note that *n* accounts for the number of slices while *N* accounts for the number of animals.

## 3. RESULTS

### 3.1. NPY concentration shows a sex- and age-dependent decrease

Several studies report a decline in NPY secretion, NPY mRNA, Y1-R, and NPY immunoreactivity during aging ^3,13,14,27^. Therefore, we investigated potential age-mediated changes in NPY concentration and Y1-R expression on the DG tissue of young-adult and aged male and female mice. In agreement with previous findings ^23^, NPY concentration in the DG was significantly lower in females than in males (F_(1,18)_=54.08, p<0.0001, two-way ANOVA; post-hoc comparison: young-adult females vs. young-adult males: p<0.0001, Fisher’s LSD test; Fig. 1B). This sex difference persisted through aging (post-hoc comparison: aged females vs. young-adult males: p<0.0001; young-adult females vs. aged males: p=0.0026; aged females vs. aged males: p=0.0002, Fisher’s LSD test; *N*=5 (young-adult males), *N*=5 (aged males); *N*=6 (young-adult females); *N*=6 (aged females); Fig. 1B). Strikingly, aged male mice showed reduced NPY levels in comparison to young-adult male mice (F_(1,18)_=5.851, p=0.0264, two-way ANOVA; post-hoc comparison p=0.0412, Fisher’s LSD test; Fig. 1B), while NPY concentration in females remained unchanged during aging (post-hoc comparison young-adult vs. aged anestrus females: p=0.2537; Fig. 1B). No age-dependent changes in Y1-R expression were evident (sex: F_(1,19)_=0.1691, p=0.6855; age: F_(1,19)_=0.07256, p=0.7906; interaction: F_(1,19)_=0.8460, p=0.3692; two-way ANOVA, *n*=5 (young-adult males); *N*=5 (aged males); *N*=7 (young-adult females); *N*=6 (aged females); Fig. 1C) in the DG. To summarize, only aged male mice showed reduced NPY levels in the DG compared to young-adult mice.

**Figure 1:**
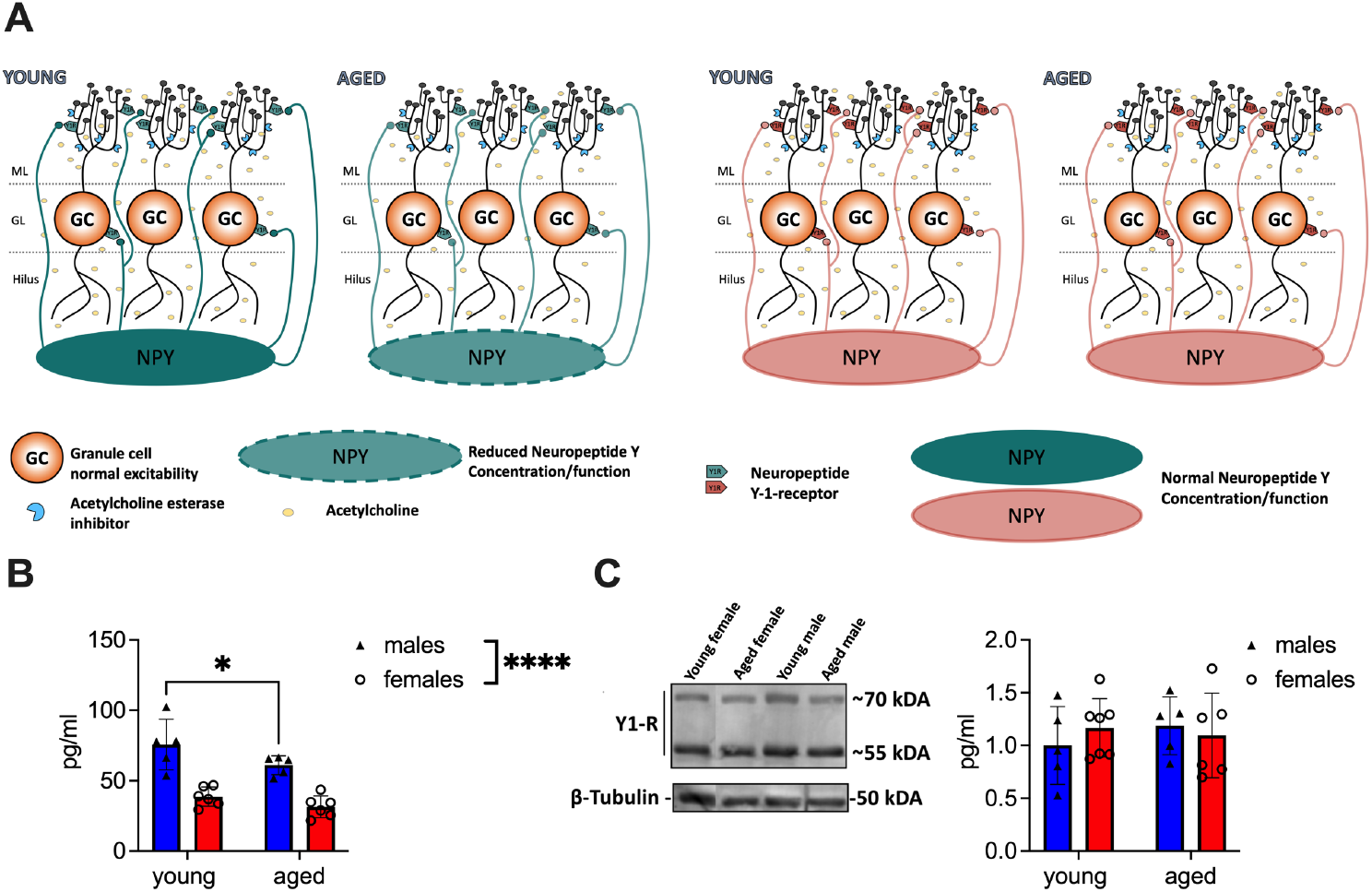
Aged male mice show reduced NPY concentration in the DG. (**A)** Scheme illustrating the reduction of NPY expression in the local circuitry. Reduced levels of NPY are evident in aged male mice, whereas Y1-R expression remains during aging. The well-established reduction of ACh is indicated with light yellow circles. **(B)** Tissue concentration of NPY in the DG decreases in males during aging. Female mice express low but unaltered NPY levels. **(C)** Y1-R expression measured by immunoblotting does not differ between sexes or over lifetime. A representative WB membrane with one example of each sex and age is presented. ACh, acetylcholine. Changes in NPY concentration (group comparison): *, p < 0.05, ****, p < 0.0001.

### 3.2. LTP induction at the MPP-DG synapse shows sex differences in an age-dependent manner

Since a decline of the NPYergic system has been related to memory impairment ^28^ we evaluated LTP at the MPP-DG synapse as the cellular correlate of memory ^29^ in both sexes and ages. Four repetitions of a TBS train were applied to reliably induce LTP under intact inhibition in young-adult males (t_(8)_ =2.056, p=0.0369, paired t-test, one-tailed, *n*=9; Fig. 2B) and females (p=0.0186, Wilcoxon test, one-tailed, *n*=10; Fig. 2B’), shown by an increased fEPSP slope. A general impairment of LTP induction and maintenance in the DG *in vitro* of middle-aged rodents has previously been shown by Schreurs and colleagues (2017) using micro-array recordings ^30^, but changes of synaptic plasticity properties of aged female mice in the dDG have so far not been investigated. Our data confirmed that LTP is abolished in aged male mice under intact inhibition (t_(9)_=0.9207, p=0.1906, paired t-test, one-tailed, *n*=10; Fig. 2B), as shown by the unchanged fEPSP slope after 4x TBS. By contrast, 4x TBS induced LTP successfully at the MPP-DG synapse of aged female mice (t_(7)_=2.019, p=0.0416, paired t-test, one-tailed, *n*=8; Fig. 2B’).

**Figure 2:**
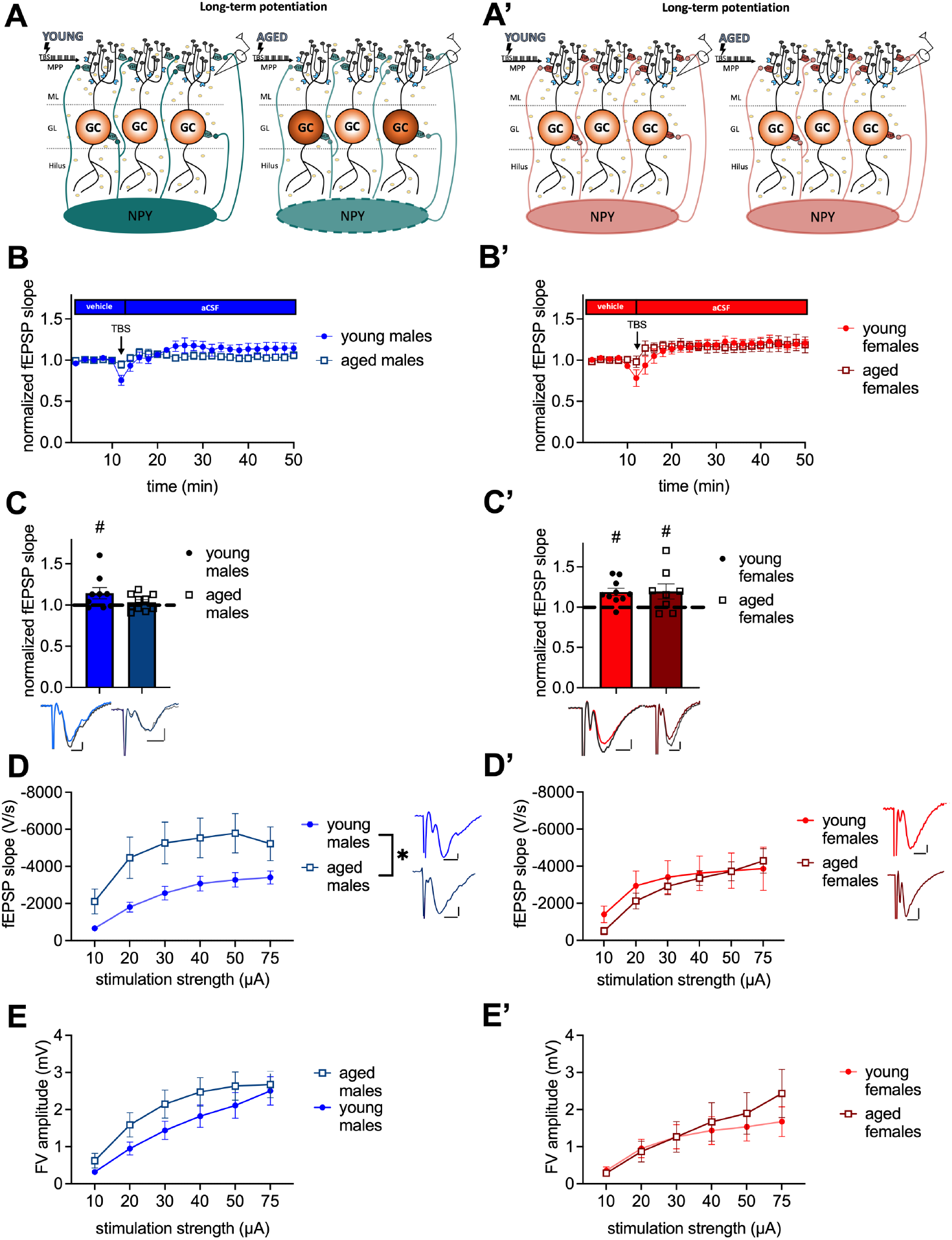
MPP-DG LTP is abolished in aged male mice, while baseline excitability is increased. **(A)** Scheme illustrating the postulated impact of reduced NPY levels in aged males on LTP and GC excitability. **(B)** Timeline of MPP-DG LTP with the average values from young-adult and aged male mice. **(C)** Significant LTP is evident in young-adult but not aged males, as seen during the last 10 min of the recording. Representative fEPSP traces are plotted beneath. (pre-TBS colored, post-TBS grey). **(D)** Post-synaptic excitability is increased in aged male mice compared to young-adult males. Representative fEPSP traces are plotted at 30 µA stimulation strength. Scale bar x-axis 2 ms each and scale bar y-axis: 1 mV each. **(E)** Pre-synaptic excitability is not significantly altered in aged males. **(A’)** Scheme illustrating the postulated impact of reduced NPY levels in aged females on LTP and granule cell excitability. **(B’)** Timeline of MPP-DG LTP with the average values from young-adult and aged female mice **(C’)** Significant LTP is evident in both young-adult and aged female mice, as seen during the last 10 min of the recording. Representative fEPSP traces are plotted beneath (pre-TBS colored, post-TBS grey). **(D’)** Post-synaptic excitability is unaltered between young-adult and aged females. Representative fEPSP traces are plotted for young-adult and aged animals at 30 µA stimulation strength. Scale bar x-axis: 2 ms each and y-axis: 1 mV. **(E’)** Pre-synaptic excitability is not significantly altered in aged females. DMSO, dimethylsulfoxid. Age effect (group comparison): *, p< 0.05; MPP-DG LTP induction (within slice comparison): #, p < 0.05.

Potential aging-related changes in circuit functions were further assessed by evaluating post-synaptic and pre-synaptic excitability. Analysis of I-O curves revealed an increase in post-synaptic excitability in aged male mice in comparison to young-adult male mice (age: F_(1, 35)_=4.243, p=0.0469; interaction: F_(5, 175)_=3.032, p=0.0119, repeated-2-way ANOVA; Fig. 2D). The post-hoc comparison showed that excitability was increased at 10, 20, 30, 40, and 50 µA (post-hoc comparison: 10 µA: p=0.0476; 20 µA: p=0.0302; 30 µA: p=0.0313; 40 µA: p=0.0405; and 50 µA: p=0.0360, Fisher’s LSD, *n*=20 (aged) and n=17 (young-adult); Fig. 2D). No change was observed in the pre-synaptic fiber volley (FV) amplitude of the aged male mice (F_(1, 34)_=1.377, p=0.2487, repeated 2-way ANOVA, *n*=20 (aged) and n=16 (young-adult); Fig. 2E). In contrast to males, post-synaptic excitability was unaltered in aged females compared to young-adult females (age: F_(1, 15)_=0.1063, p=0.7489, interaction: F_(5, 75)_=1.261, p=0.2899, repeated two-way ANOVA; *n*=10 (young-adult females); *n*=7 (aged females); Fig. 2D’). Furthermore, the pre-synaptic excitability was unchanged in the aged females (age: F_(1, 14)_=0.1623, p=0.6932, interaction: F_(5, 70)_=1.866, p=0.1114, repeated two-way ANOVA; *n*=9 (young-adult females) *n*=7 (aged females); Fig. 2E’). In summary, LTP in aged male mice is abolished, while it persists in aged females. Age-meditated LTP sex differences are accompanied by increased post-synaptic excitability in males, while post-synaptic excitability is unchanged during aging in females.

### 3.3. Cholinergic activation rescues MPP-DG LTP of aged male mice in a Y1-R-dependent manner

Based on our biochemical and physiological findings the potential involvement of the NPYergic system in aging-related synaptic functionality changes was further investigated in male mice.

The beneficial effect of AChE inhibitor for AD treatment and on CA1 LTP in aged rodents is well documented ^18^. By contrast, the beneficial effect of the cholinergic treatment on MPP-DG LTP and the involved mechanism have not been fully resolved yet. Based on our laboratory’s investigation that the ACh-mediated release of NPY is essential for memory retrieval in the DG of young-adult male mice ^7^, we evaluated the importance of Y1-R activation after cholinergic activation on MPP-DG LTP by applying the AChE inhibitor physostigmine (PHY 2µM) and the combination of PHY and the selective Y1-R antagonist BIBP3226 (1µM). Under cholinergic activation in young-adult male mice, LTP was facilitated compared to that in controls, regardless of Y1-R blockage (control: t_(7)_=2.469, p=0.0214, paired t-test, one-tailed, *n*=8; PHY: t_(8)_=2.403, p=0.0215, paired t-test, one-tailed, *n*=9; PHY + BIBP3226: t_(9)_=2.265, p=0.0249, paired t-test, one-tailed, *n*=10; Fig. 3D). Furthermore, cholinergic activation by PHY rescued the LTP in aged male mice (control: t_(7)_=1.608, p=0.0645, Wilcoxon test, one-tailed, *n*=9; PHY: t_(12)_=3.821, p=0.0012, paired t-test, one-tailed, *n*=13; Fig. 3D). However, this rescue was dependent on the activation of the Y1-R as shown by its abolition upon additional BIBP3226 application (t_(10)_=1.569, p=0.0739, paired t-test, one-tailed, *n*=11; Fig. 3D). A treatment effect was depicted (F_(2, 53)_=6.338, p=0.0034, two-way ANOVA; Fig. 3D) by the increased LTP strength after PHY application in young-adult males (p=0.0300, Fisher’s LSD test; Fig. 3D) also compared to the LTP in aged male mice (control: p=0.0085; PHY+ BIBP3226: p=0.0022, Fischer’s LSD test; Fig. 3E). Also, in aged mice LTP strength was enhanced by PHY compared to control (p=0.0500, Fisher’s LSD test; Fig 3D) and significantly decreased after additional Y1-R blockade (p=0.0158, Fisher’s LSD test; Fig. 3D).

**Figure 1:**
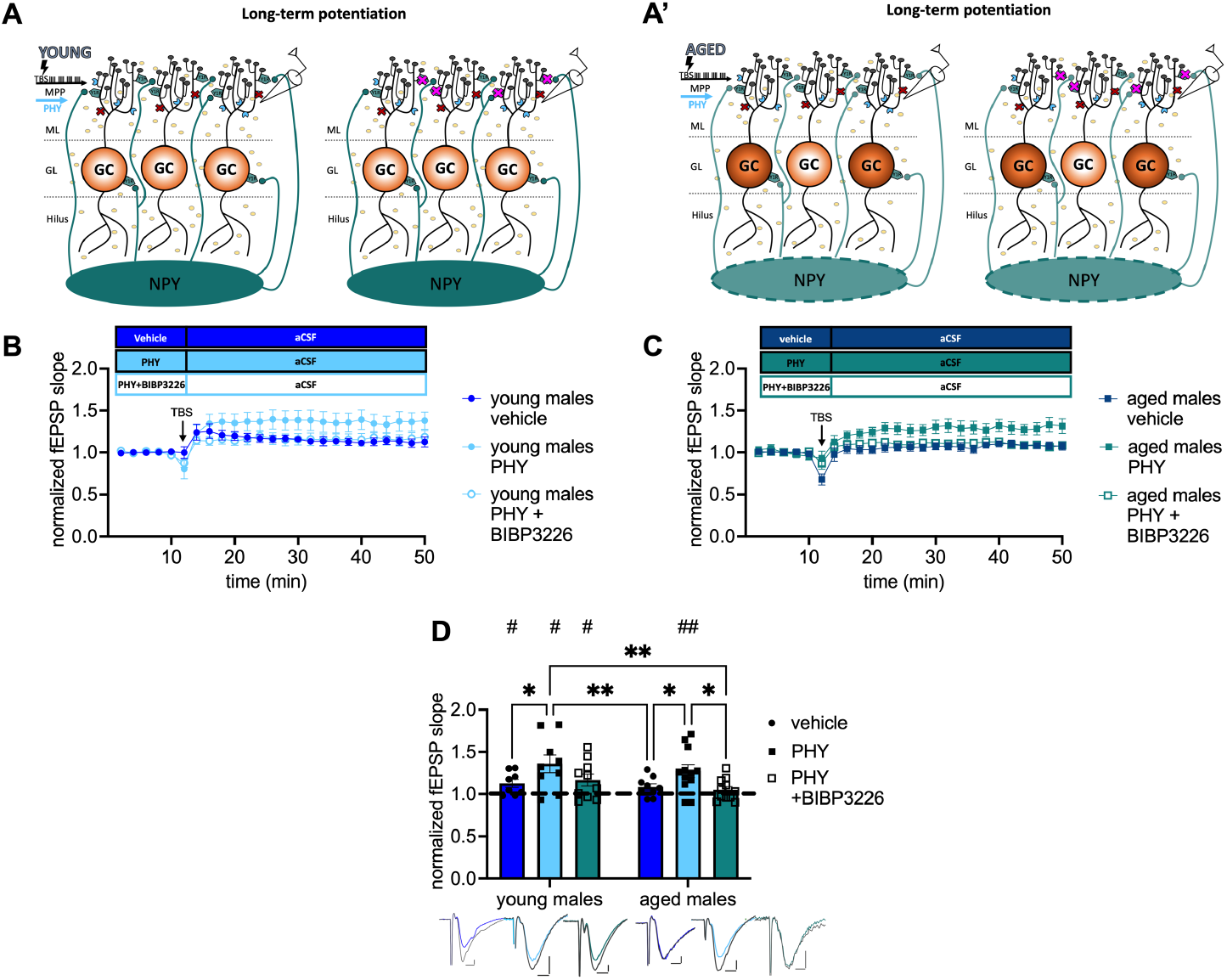
Cholinergic rescue of MPP-DG LTP in aged male mice is Y1-R-dependent. **(A, A’)** Scheme illustrating the postulated involvement of NPY release in the cholinergic stimulation-induced rescue of LTP in young-adult **(A)** and aged **(A’)** male mice. **(B)** Timeline of recosrdings from young-adult males with and without physostigmine (PHY) and Y1-R antagonist BIBP3226. **(C)** Corresponding timeline from aged males. **(D)** Physostigmine enhances LTP in young-adult males and reinstates it in aged males. In aged males, LTP is entirely prevented by the co-administration of BIBP3226. Representative fEPSP traces are plotted beneath (pre-TBS colored, post-TBS grey). Scale bar x-axis: 2 ms each and y-axis: 1mV (young-adult males) and 0.4 mV (aged males). Changes in LTP strength (group comparison): *, p < 0.05, **, p < 0.01; LTP induction (within slice comparison): #, p < 0.05, ##, p < 0.01.

To note, blockade of Y1-R alone did not affect MPP-DG LTP induction in young-adult nor in aged animals (see supplements Fig. A.1C). Nevertheless, Y1-R blockade alone led to significant differences in LTP strength between the groups (see supplements Fig. A.1C). These results indicate a crucial role of Y1-R activation under moderate cholinergic activation in the MPP-DG LTP of aged male mice.

### 3.4. Decrease of baseline transmission upon pharmacological cholinergic activation depends on Y1-R activation in young-adult but not in aged male mice

Next, we tested the impact of moderate cholinergic activation on the Y1-R activation during neurotransmission. Strikingly, treatment (F_(2, 72)_=8.556 p=0.0005, two-way ANOVA; Fig. 4E) and age effects (F_(1, 72)_=5.918 p=0.0175, two-way ANOVA; Fig. 4E) were evident. Baseline transmission decreased after PHY application in young-adult (post-hoc comparison to young-adult males: p=0.0186, Fisher’s LSD test, *n*=10 (young-adult males), *n*=14 (young-adult males + PHY); Fig. 4D) and in aged male mice (post-hoc comparison to aged males: p=0.0068, Fisher’s LSD test, *n*=14 (aged males), *n*=13 (aged males + PHY); Fig. 4D). Strikingly, additional blockade of the Y1-R in young-adult male mice increased neurotransmission back to baseline (post-hoc comparison to: young-adult male mice + PHY: p=0.0009; aged male mice + PHY: p<0.000; *n*=16 (young-adult males + PHY + BIBP3226), *n*=13 (aged males + PHY); Fig. 4D). However, this reversal was not observed in the aged males, as evident from the unchanged fEPSP slope (post-hoc comparison to aged males mice + PHY: p=0.1353, Fisher’s LSD test, *n*=11 (aged males + PHY + BIBP3226); *n*=13 (aged males + PHY); Fig. 4D) which was also not significant when compared to young-adult male mice + PHY (post-hoc comparison: p=0.5316, Fisher’s LSD test, *n*=11 (aged males PHY + BIBP3226); *n*=14 (young-adult males + PHY); Fig. 4D). Moreover, fEPSP slope remained lower than in young-adult male mice + PHY + BIBP3226 (post-hoc comparison: p=0.0113, Fisher’s LSD test, *n*=11 (aged males PHY + BIBP3226); *n*=16 (young-adult males + PHY + BIBP3226); Fig. 4D). An interaction (F_(2, 72)_=1.023 p=0.3642, two-way ANOVA; Fig. 4E) was not evident. To note, MPP-DG neurotransmission remained unchanged after selective Y1-R blockade independent of age, evident by the unchanged fEPSP slope (see supplements Fig. A.2D). Nevertheless, an interaction effect reveals a significant increase of fEPSP slope in young-adult males treated with BIBP3226 compared to the corresponding aged group (see supplements Fig. A.2D) Taken together, increased cholinergic tonus mediated a decrease of MPP-DG neurotransmission in male mice in both young-adult and aged male mice. In contrast, the reversal of by Y1-R blockade was age-dependent and moderate cholinergic activation mediated Y1-R activation specifically in young-adult animals.

**Figure 4:**
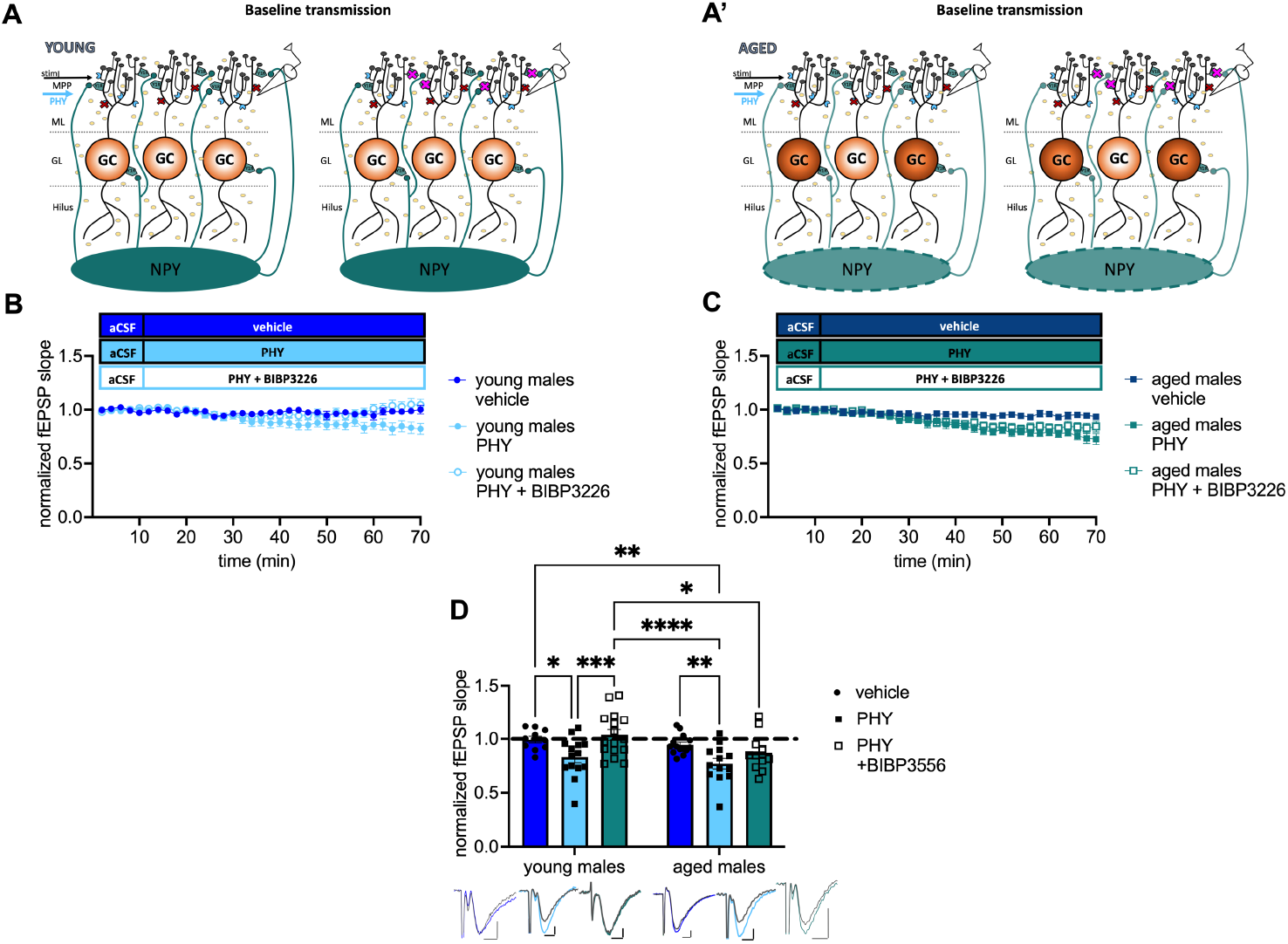
Pharmacological cholinergic activation promotes inhibition of excitability in a Y1-R and age-dependent manner. **(A, A’)** Scheme illustrating the postulated involvement of NPY release in the cholinergic stimulation-induced decrease in baseline transmission. **(B)** Timeline of baseline transmission recordings from young-adult males with and without PHY and the Y1 antagonist BIBP3226. **(C)** Corresponding timeline from aged males. **(D)** PHY decreases baseline transmission in a Y1-R-dependent manner in young-adult but not aged males. Representative fEPSP traces are plotted beneath. Scale bar x-axis: 2 ms each and y-axis: 1 mV (young-adult mice control); 0.5 mV (aged mice PHY + BIBP3226) and 0.4 mV (young-adult and aged mice PHY, PHY + BIBP3226, and aged mice control; pre-application colored, post-application grey). Changes in fEPSP slope after Y1-R blockade (group comparison) *, p < 0.05, **, p < 0.01, ***, p < 0.001, ****, p < 0.0001.

### 3.5. Application of NPY rescues LTP in aged male mice

The data above indicate a crucial role of Y1-R activation to maintain plasticity during aging. Furthermore, Santos and colleagues conducted a beneficial effect of intracerebroventricular NPY infusion on spatial memory in young male mice ^31^. Therefore, the potential impact of NPY (1 µM) on plasticity in aged male mice was investigated. Application of NPY rescued LTP (t_(10)_=3.222, p=0.0046, paired t-test, one-tailed, *n*=11; Fig. 5C) as evidenced by the increased fEPSP slope after TBS under NPY. Furthermore, NPY application led to a significantly increased LTP compared to vehicle controls (p=0.0087, Mann-Whitney test, two-tailed, *n*=11 (NPY); *n*=5 (control); Fig. 5C). By contrast, baseline neurotransmission was unchanged after NPY application (t_(14)_=0.5243, p=0.6083, unpaired t-test, two-tailed, *n*=5 (control), *n*=11 (NPY); Fig. 5E). This experiment underlines the importance of NPYergic tonus for successful LTP induction in aged male mice.

**Figure 5:**
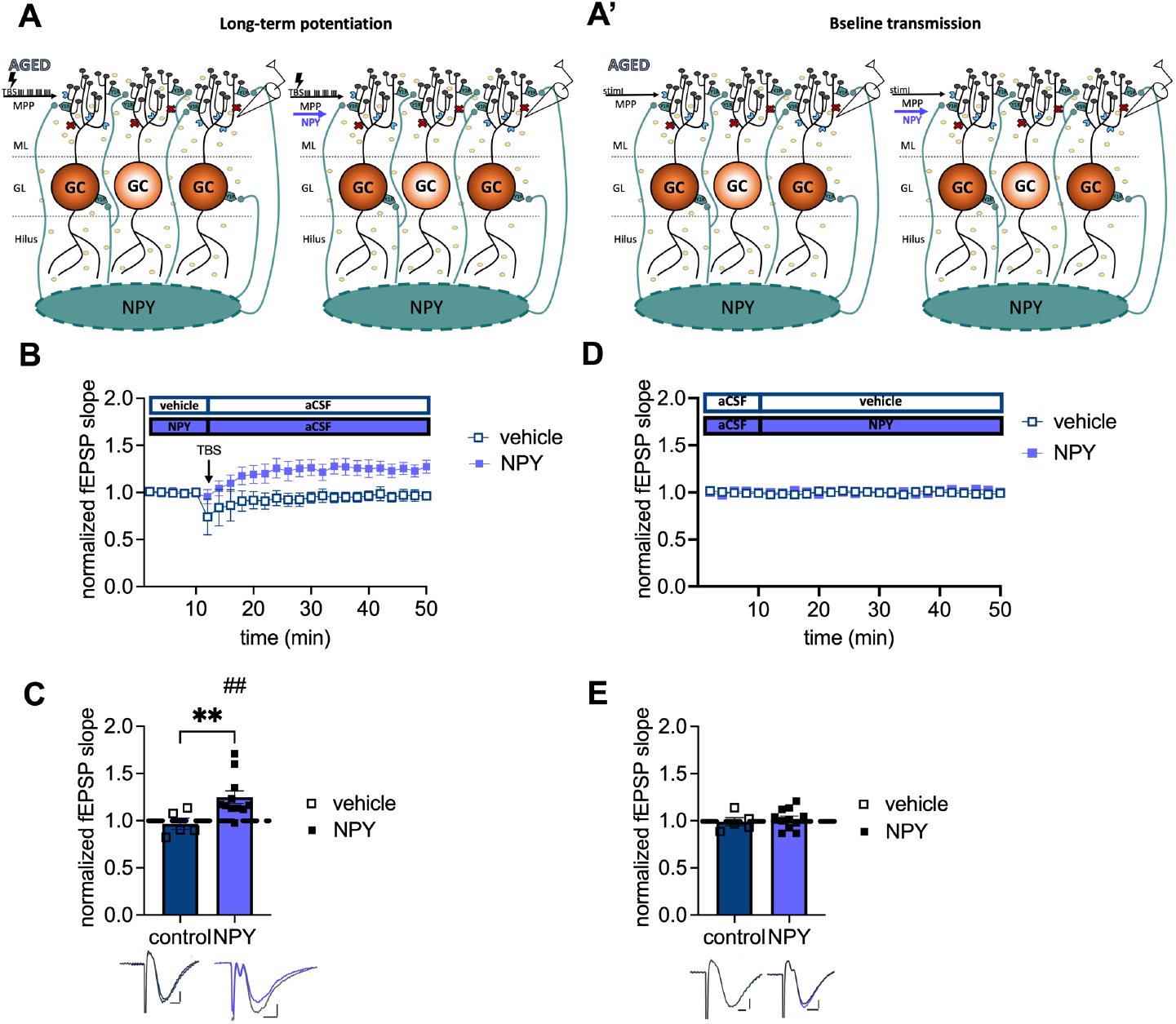
Increased NPYergic neurotransmission rescues LTP in aged male mice. **(A)** Scheme illustrating the impact of increased NPY levels mediated by exogenous NPY application on plasticity **(A)** and GC excitability **(A’)** in aged male mice. **(B)** Timeline of plasticity recordings from aged males with and without exogenous NPY. **(C)** NPY reinstates a robust LTP in aged males. **(D)** Timeline of baseline transmission recordings from aged males with and without exogenous NPY. **(E)** Exogenous NPY application does not impact neurotransmission in aged male mice. Representative fEPSP traces are plotted beneath (pre-TBS/drug application colored and post-TBS/after drug application grey). Scale bar x-axis: 2 ms each and scale bar y-axis: 0.4 mV (control LTP) and 1 mV (transmission and NPY LTP). DMSO, dimethylsulfoxide. MPP-DG LTP strength (group comparison): **, p < 0.01 and MPP-dDG LTP induction (in slice comparison): ##, p < 0.01.

## 4. DISCUSSION

In this study, we demonstrate a critical role of NPY in the loss of synaptic plasticity in the aged mouse DG and its recovery through stimulation of cholinergic neuromodulation. These data are of particular relevance considering the reported loss of NPYergic interneurons in the aged DG and reflected in aging-induced changes in NPY content of the structure, specifically in male mice.

To investigate the involvement of NPYergic interneurons in age-related changes of DG plasticity, we established in slice preparation protocols for induction of LTP at the MPP-DG synapse under intact inhibition. These protocols produced a mild but reliable potentiation that are comparable to previous studies under intact GABAergic tonus ^25,32^. Thereby we were able to demonstrate the expected loss of LTP in aged male mice and its recovery upon cholinergic stimulation, which is in line with various in vivo studies reporting impaired LTP at the PP-DG synapse of aged male rodents ^33,34^. By contrast we found that LTP was maintained in the DG of naturally aged females even in the absence of an AChE inhibitor, in line with findings in a female Alzheimer’s diseases mouse model ^35^.

Given the high but age-dependent decaying expression of NPY in the male DG and the constant low levels of this peptide in females, we tested a possible involvement of the peptide in the observed plasticity effects of male mice. In fact, we observed that recovery of LTP upon cholinergic stimulation in aged animals was dependent on NPY-Y1 receptors, the major post-synaptic NPY receptor type in the DG (Fig. 3E). By contrast, the reduction of baseline transmission induced by cholinergic stimulation persisted under NPY-Y1 receptor blockage in the aged animals, in contrast to young (Fig. 3E). Moreover, unchanged pre-synaptic response in aged mice (Fig. 2E) indicating that MPP-DG LTP deficits were related to local changes in the DG during aging rather than alterations of baseline excitability or a non-functional input from the EC ^33,36^.

A decline of the projections from the MS to the DG during aging has been previously described _15_. These cholinergic projections from the MS target both GCs and hilar interneurons, including NPY/SST-containing HIPP cells and parvalbuminergic basket cells, and axo-axonic cells ^37^. HIPP cells are vulnerable to a cholinergic decline ^27^ and mediate cholinergic-induced NPY-Y1 receptor stimulation at the DG GCs ^7^. Non-fast spiking interneurons, including HIPP cells, are strongly recruited after strong PP activation to maintain the quiescence of GC ^38^ and NPY is known to reduce glutamatergic transmission and the release of Ca^2+^ in GCs ^39^. Moreover, a reconfiguration effect of cholinergic transmission on the hilar interneuron network, particularly SST^+^ (and putatively NPY^+^) cells and PV+ basket cells, has been reported ^37^ that might mediate the facilitatory effect of muscarinic transmission on LTP at MPP-DG synapses ^37,40^. Intriguingly, a subpopulation of PV+ basket cells and axo-axonic cells in the hilus also expresses NPY ^41^. It is thus conceivable that enhanced cholinergic tonus might act in part by stimulating NPY release from residual hilar interneurons in the aged animals to overcome the aging-related local deficit in this peptide (Fig. 1B). In support of this explanation, we could show that application of NPY alone, in the absence of acetylcholinesterase blockade was sufficient to reinstate the ability for induction of LTP (Fig. 5C). Interestingly, both expression and physiological data indicate that NPY appears to play a sex-specific role in DG function during aging, more relevant to male than to female mice.

We considered the possibility that the effect of NPY may at least in part be mediated by recovering excitation-inhibition balance in the DG, as prominent post-synaptic hyperexcitability could be observed in the aged animals (Fig. 2D). E-I balance is essential for maintaining signal-to-noise ratio, information capacity, and sparse GC activation in the DG _42,43_. Aged rodent DG displays increased glutamate release ^44,45^, a higher population spike to a given PP stimulus, and a reduced voltage threshold ^4^. Moreover, increased E-I ratio in aged rats after LPP stimulation correlates with reduced cognitive performance ^46^ and enhanced post-synaptic inhibitory strength has been found in cognitively unimpaired-aged rats ^47^. NPY Y1-receptor activation might be able to reduce glutamate release and Ca^2+^ influx and, with that, reduce GC excitability ^39^. However, the application of NPY in our experiments did not affect MPP-DG transmission (Fig. 5E), and NPY-Y1 receptor blockage failed to interfere with the cholinergic-induced reduction of excitatory inhibition in aged mice (Fig. 4D). This is in line with previous studies in young-adult animals showing no or minor effects of NPY on PP-evoked EPSPs in the molecular layer ^39^. Thus, in aged mice, NPYergic transmission appears to become particularly relevant for the induction of plasticity in the MPP-DG pathway.

Various studies have demonstrated a role of NPY and NPY Y1 receptors in emotional and learning deficits in Alzheimer’s disease models and their cholinergic recovery ^31,48^. However, a memory-enhancing effect of NPY overexpression in the hippocampus and hypothalamus of rats is not observed in middle-aged animals, while its anxiolytic-like effects persist ^49^. Given the prominent role of the DG interneurons in both cognition and emotion ^50–52^, NPY appears to act as a network modulator that integrates, in a sex-dependent manner, the response of the hilar network to stress- and aging-induced alterations in excitability and plasticity and may lend itself as a target of intervention.

## 5. ABBREVIATIONS

aCSF: artificial cerebrospinal fluid
AChE: acetylcholine esterase
DG: dentate gyrus
fEPSP: field excitatory post-synaptic potential
GC: granule cells
LTP: Long-term potentiation
ML: molecular layer
MPP: medial perforant path
NPY: Neuropeptide Y
PHY: physostigmine
PP: perforant path
SST: somatostatin
TBS: theta-burst stimulation
Y1-R: neuropeptide Y type 1 receptor

## 6. ACKNOWLEDGEMENTS

We are grateful to F. Webers, S. Stork, and A. Koffi von Hoff for expert technical assistance and A. Bohnstedt, D. Al-Chakmakchi, and M. Meier for excellent animal care.

## 7. AUTHOR CONTRIBUTIONS

Conceptualization, KK, G.Ç., and OS; Methodology, KK, G.Ç., and OS Investigation, KK and M.DA.; Writing – Original draft, KK; Writing – Review & Editing KK, G.Ç., and OS; Funding Acquisition, OS; Resources, OS; Supervision, G.Ç. and OS.

