## Supplement Figures A1 and A2 for "Increasing NPYergic transmission in the hippocampus rescues aging-related deficits of long-term potentiation in the mouse dentate gyrus"

**SUPPLEMENTS**

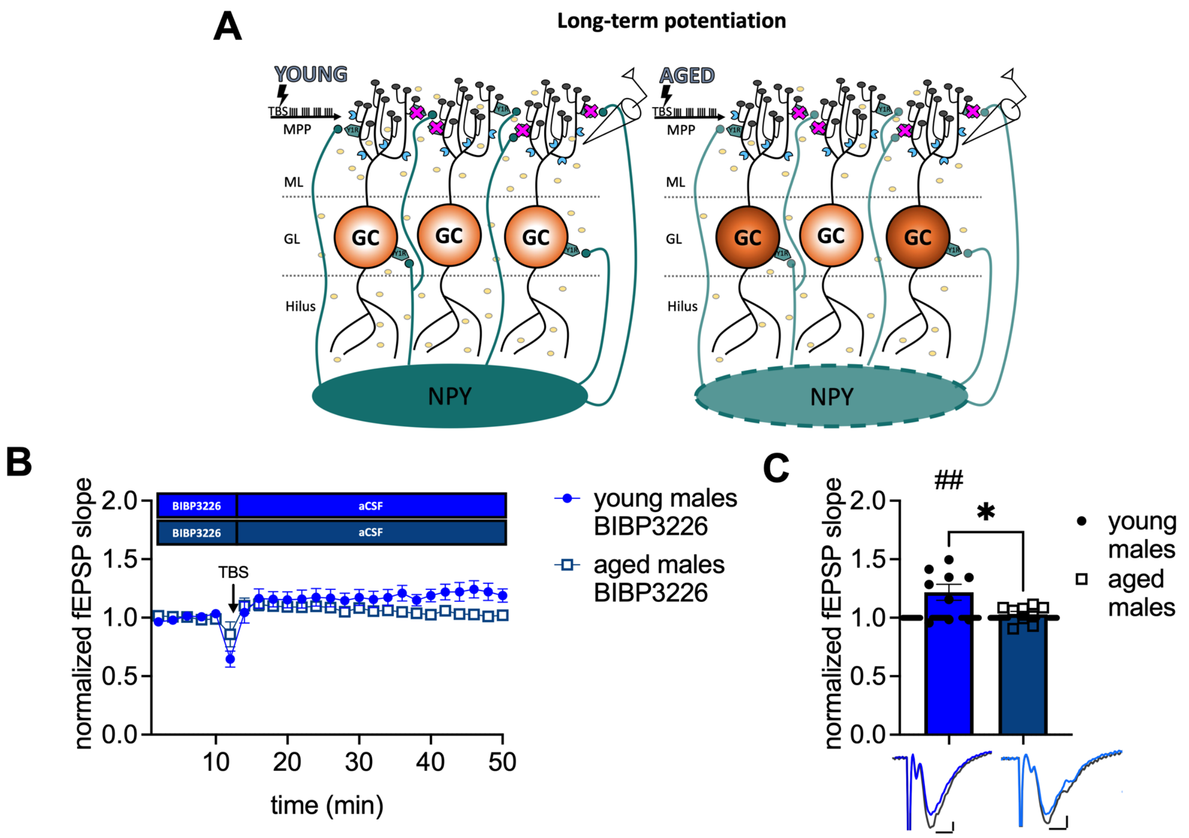


**Figure A.1: Y1-R blocking does not facilitate MPP-DG LTP. (A)** Scheme illustrating the postulated impact of reduced NPY concentration on LTP. **(B)** Timeline of recordings from young-adult and aged males with the Y1-R antagonist BIBP3226. **(C)** BIBP3226 does not impact LTP in male mice. LTP induction is successful under Y1-R blockade in young-adult animals (blue; $t_{(8)}$=3.080, p=0.0076, paired t-test, one-tailed, n=9) but not in aged male mice (dark blue; $t_{(8)}$=1.044, p=0.3271, paired t-test, one-tailed, n=9). Nevertheless, a difference in LTP strength between young-adult and aged males is demonstrated ($t_{(16)}$=2.591, p=0.0197, unpaired t-test, two-tailed). Representative fEPSP traces are plotted below (pre-TBS colored; post-TBS grey). Scale bar x-axis: 2 ms each and y-axis: 1 mV (young males) and 0.4 mV (aged males). DMSO = dimethylsulfoxid. MPP-DG LTP strength (group comparison): *, $p < 0.05$; MPP-dDG LTP induction (in slice comparison): ##, $p < 0.01$.

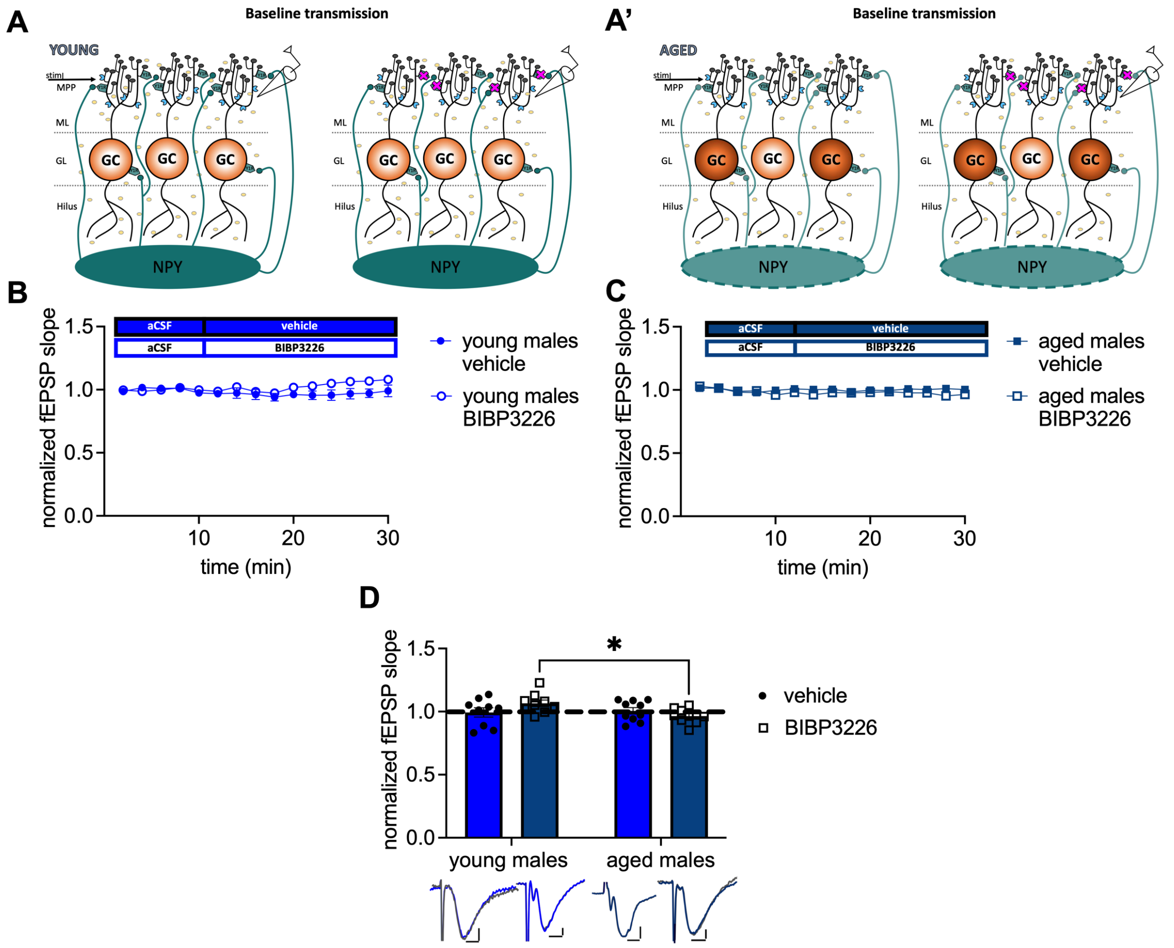


**Figure A.2: Y1-R blockade has no effect on DG neurotransmission in both ages. (A)** Scheme illustrating the postulated impact of reduced NPY concentration on baseline transmission. **(B)** Timeline of baseline recordings from young-adult males with the Y1-R antagonist BIBP3226. **(C)** Corresponding timeline from aged males. **(D)** BIBP3226 impact neurotransmission in male mice ($F_{(1, 33)}$=4.187 p=0.0488, two-way ANOVA, n=28 (young-adult males + BIBP3226); n=29 (young-adult males control); n=9 (aged males + BIBP3226); n=10 (aged males control). However, baseline transmission is unchanged between young-adult male mice (post-hoc comparison: p=0.0777, Fisher's LSD test) and aged males (p=0.2953, Fisher's LSD test) but stays lower in aged male mice compared to young-adult male mice upon BIBP3226 application (p=0.0159, Fisher's LSD test). Representative fEPSP traces are plotted below. Baseline is colored and after drug application in grey. Scale bar x-axis: 2 ms each and y-axis: 0.4 mV each. DMSO = dimethylsulfoxide. MPP-DG transmission (group comparison): *, $p < 0.05$.
